# Computational toolbox for ultrastructural quantitative analysis of filament networks in cryo-ET data

**DOI:** 10.1101/2021.05.25.445599

**Authors:** Georgi Dimchev, Behnam Amiri, Florian Fäßler, Martin Falcke, Florian KM Schur

**Author notes:** Equal contribution (Georgi Dimchev, Behnam Amiri).

## Abstract

A precise quantitative description of the ultrastructural characteristics underlying biological mechanisms is often key to their understanding. This is particularly true for dynamic extra- and intracellular filamentous assemblies, playing a role in cell motility, cell integrity, cytokinesis, tissue formation and maintenance. For example, genetic manipulation or modulation of actin regulatory proteins frequently manifests in changes of the morphology, dynamics, and ultrastructural architecture of actin filament-rich cell peripheral structures, such as lamellipodia or filopodia. However, the observed ultrastructural effects often remain subtle and require sufficiently large datasets for appropriate quantitative analysis. The acquisition of such large datasets has been enabled by recent advances in high-throughput cryo-electron tomography (cryo-ET) methods. However, this also necessitates the development of complementary approaches to maximize the extraction of relevant biological information. We have developed a computational toolbox for the semi-automatic quantification of filamentous networks from cryo-ET datasets to facilitate the analysis and cross-comparison of multiple experimental conditions. GUI-based components simplify the manipulation of data and allow users to obtain a large number of ultrastructural parameters describing filamentous assemblies. We demonstrate the feasibility of this workflow by analyzing cryo-ET data of untreated and chemically perturbed branched actin filament networks and that of parallel actin filament arrays. In principle, the computational toolbox presented here is applicable for data analysis comprising any type of filaments in regular (i.e. parallel) or random arrangement. We show that it can ease the identification of key differences between experimental groups and facilitate the in-depth analysis of ultrastructural data in a time-efficient manner.

## Introduction

Cryo-electron tomography (cryo-ET) provides high-resolution insights into natively preserved biological environments in cells and tissues. Beyond its use for *in situ* structure determination (*1, 2*), its main strength lies in its ability to provide contextual information for the molecules under study, such as the higher-order arrangement of proteins in cells. This information can be linked to functional data to provide a holistic quantitative description of cellular processes. In this regard, cryo-ET with its resolution on the level of individual molecules is well positioned to complement experimental data obtained by other modalities, such as genetic perturbation experiments or light-microscopy imaging.

One major challenge in cryo-ET is the extraction of statistically relevant quantitative parameters from sufficiently large datasets. Several inherent attributes of the method impede large-scale analysis, including the low signal to noise (SNR) ratio in tomograms, the complexity of cellular data, and the need of appropriate computational tools to extract meaningful biological data. Hence, while the potential of cryo-ET as a qualitative method is commonly accepted for applications where the analysis of a few tomograms is sufficient to detect and describe novel subcellular features, its potential as a quantitative technique to compare subtle differences among genetically distinct samples is not yet fully realized.

Recent improvements in cryo-EM sample preparation (*3*), automated EM data acquisition (*4*–*6*), image processing workflows (*7*), and data analysis allow the evaluation of large datasets and comparison of various *in situ* features between multiple experimental conditions. These improvements, although very suitable for being combined with the nowadays relatively straightforward genetic manipulation of cell lines via CRISPR/Cas9 techniques, are yet to be routinely applied in workflows that facilitate the high-throughput analysis and comparison of ultrastructural characteristics between genetically modified cell lines.

Studying such large datasets is a prerequisite to compensate for random errors that can occur when segmenting and vectorizing objects in tomograms. Thus, the accuracy of the obtained data ultimately depends on the quality of the tomograms and the dataset size, where the latter can compensate for errors that are tomogram-specific (i.e. caused by local variations in tomogram quality).

Characterization of molecular machineries underlying cell migration strongly benefits from quantitative descriptions. This is particularly true for the actin cytoskeleton and its associated regulatory proteins (*8*). Together, they form dynamic higher-order structures at the leading edge of migrating cells including sheet- or finger-like protrusions, such as lamellipodia, and microspikes or filopodia. The ultrastructural and morphological characterization of these assemblies in wild type or genetically modified cells, combined with experiments elucidating cellular dynamics, can provide an accurate description of the role of selected players in the initiation and maintenance of actin networks or how actin filaments produce forces in a variety of cellular mechanisms (*9*– *11*).

(Cryo-) electron tomography has provided ultrastructural insights into distinct actin filament assemblies, such as lamellipodia, filopodia, actin waves or pathogen-mediated filament networks (*11*– *17*). Specifically, major progress was achieved by introducing computational tools to vectorize filaments, either based on template matching or using the localized radon transform, to then derive parameters for entire filament networks (*15, 18*). Due to the experimental complexity, previous studies analyzed datasets ranging from a few to ∼30 tomograms (*12, 16, 19*–*21*), and the subsequent quantitation of the vectorized filament information employed single-function customized scripts predominantly to derive a limited number of parameters. However, given the ongoing developments in the cryo-ET field, theoretically, datasets with hundreds of tomograms can be acquired within a few days. An exhaustive quantitative analysis could reveal more detailed descriptions of the mechanisms underlying actin network assembly and maintenance, but requires facilitated analysis workflows that are also more easily applicable to the growing base of researchers using cryo-ET approaches.

We have developed a MATLAB-based analysis toolbox that enables the semi-automatic quantification of filamentous networks from large cryo-ET datasets. It allows for pre-processing coordinate information of filaments derived from tomograms, advanced visualization of whole structures and extraction of a large number of ultrastructural parameters as either numerical values or as figures and plots. Furthermore, the toolbox facilitates cross-comparison of experimental conditions. We demonstrate the feasibility of this workflow by comparing differentially manipulated lamellipodial actin networks and parallel actin filament arrays in protruding filopodia or non-protruding microspikes.

## Results and Discussion

### A computational toolbox facilitating ultrastructural analysis of filament-rich structures

To facilitate the adoption of a more streamlined ultrastructural analysis approach of filament populations and their characteristics in cryo-electron tomograms, we designed our computational toolbox with four key aspects in mind:

### 1) Compatibility

Our toolbox is implemented to analyze vectorized filaments thus allowing the user to employ their own method of choice to generate coordinate files of filaments from cryo-ET data (Fig. 1A). Examples for such workflows are given below: Tomograms can be preprocessed prior to vectorization using tools based on Deep learning, such as YAPiC (*22*), to segment filaments and increase the SNR. Filament vectorization can then be performed using available tools based on a template matching approach (*18*), as implemented in the commercial software Amira-Avizo (Thermo Fisher Scientific), in MATLAB-scripts using the localized Radon transform (*15*) (Suppl. Fig*-*1) or via manual filament tracking (for example in IMOD). Importantly, our toolbox is blind towards prior data vectorization approaches and requires as input the extracted filament coordinate data solely in tab-delimited format, where four columns describe the filament/object identifier and the x, y and z coordinates, respectively. Such format can be easily obtained from the published vectorization software workflows and also from IMOD after manual filament tracking.

**Figure 1.**
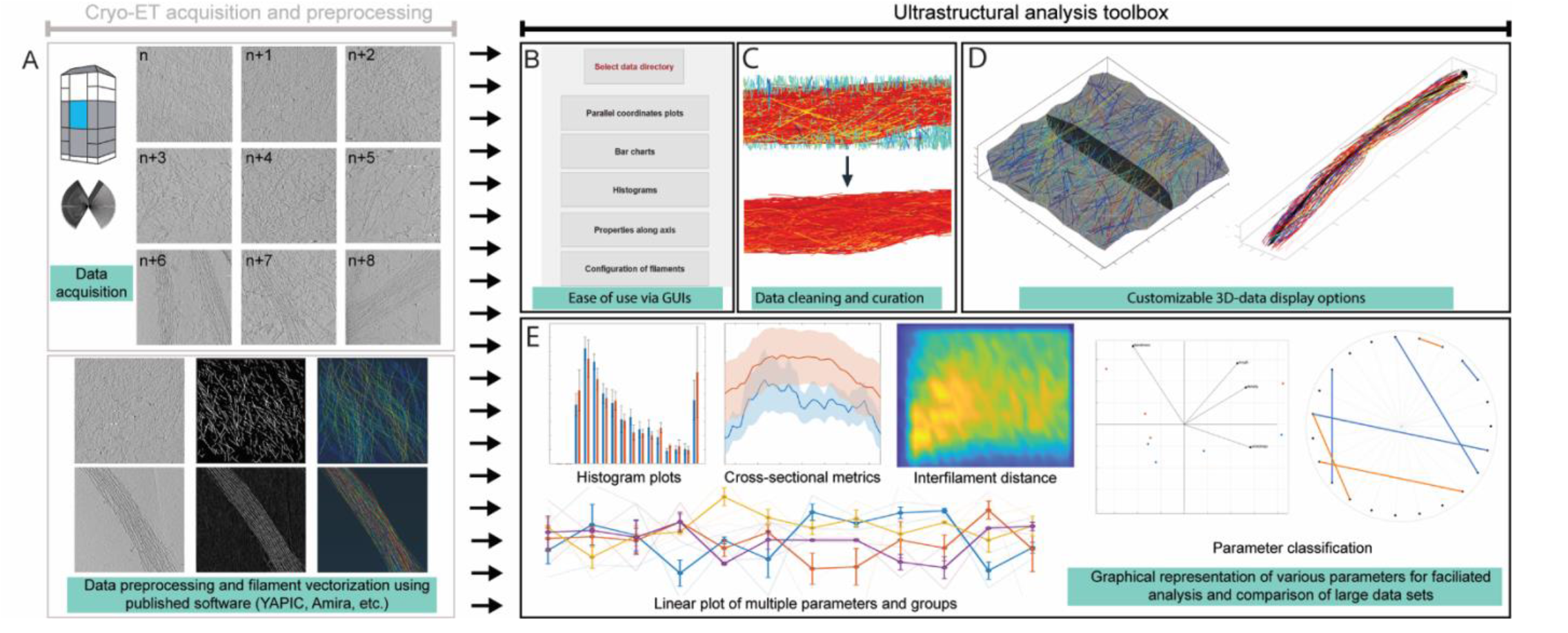
Workflow for the ultrastructural characterization of filamentous assemblies. **(A)** Tilt-series acquisition and tomogram reconstruction of datasets of variable size (upper panel) is followed by filament vectorization and extraction of filament coordinates data (lower panel). The extracted filament coordinate files can then be used as analysis input in the computational toolbox. Please note that steps described in **(A)** are not part of the presented toolbox. **(B-E)** Examples of tools and options available in the computational analysis toolbox. **(B)** GUI-based modules of the toolbox facilitate the manipulation and analysis of datasets. **(C)** Data cleaning based on user-defined ranges for filament length, bendiness or angular distribution in X/Z-axis, allows for the removal of unspecific background or exclusion of filament populations with common characteristics. **(D)** A 3D visualization module allows user-defined specific representation of sample characteristics. **(E)** Several examples of graphs and plots for the facilitated data analysis of a large number of ultrastructural characteristics.

### 2) User-friendliness & versatility

We developed our computational toolbox to require minimal MATLAB proficiency and no prior coding experience. Several graphical user interfaces (GUI) guide the user through extracting outputs from large datasets in a time efficient manner (Fig. 1B). Specifically, we have compiled the extraction of multiple predefined ultrastructural parameters from different filament architectures, such as either randomly distributed networks (e.g. lamellipodia) or quasi-parallel or bundled filaments (e.g. filopodia/microspikes) into one GUI-based step. A summary of all parameters is provided in Table 1 and 2 (see also methods section for their mathematical descriptions). These customized parameters describe whole structural features, filament ultrastructural characteristics, as well as physical properties. An exhaustive documentation file and test data is provided with the toolbox and guides the user through the individual steps and provides in-depth details on their use.

**Table 1.** Description of parameters included in the computational toolbox, specific to either quasi-parallel filament arrays or dendritic networks.

| Parameter set describing entire quasi-parallel filament structures<br>(e.g. filopodia/microspikes) |  |  |
| --- | --- | --- |
| Parameter name | Parameter short description and biological relevance | Unit<br>[value ranges] |
| Length (of structure) | The axis length of the filopodium/microspike. | nm |
| Bendiness (of structure) | Bendiness is defined as the ratio of the total axis length and its end-to-end distance. Bendiness of 1 defines a straight line. The more bent the structure, the higher the value. | a.u./ratio<br>[≥1] |
| Bending energy density (of structure) | A metric of curvature and bending property of a contour, such as the axis of a filopodium/microspike. Bent structures have a higher value of bending energy. As compared to the bendiness parameter (described above), higher values of bending energy can also reflect various contour anomalies, such as edges or sharp change in directionality/curvature of a filament. *For a more detailed mathematical description, see methods section.* | nm<sup>-2</sup> |
| Cross-sectional circularity (of structure) | Mean cross-sectional circularity of the structure averaged across 50 equidistant cross sections along the axis. Circularity is defined as the ratio of the cross-sectional area to the area of a circle with the same perimeter. It characterizes how similar the average cross section of the structure is to a perfect circle. For a perfect circular cross section circularity equals 1, while more flattened structures will have a lower value of this parameter. *For a more detailed mathematical description, see methods section.* | a.u./ratio<br>[≤1] |
| Vertical bending stiffness (of structure) | The moment of inertia of filaments in a cross section with respect to the y-axis. The parameter is associated with stiffness and describes the resistance of the structure against bending in z direction (e.g. filopodium tip rises from the substrate). Note that the parameter does not consider potential binding events between individual filaments or between filaments and other proteins, which might occur *in situ*. *For a more detailed mathematical description, see methods section.* | nm<sup>2</sup> |
| Lateral bending stiffness (of structure) | The moment of inertia of the filaments in a cross section with respect to z-axis. The parameter is associated with stiffness and describes the resistance against lateral bending, e.g. how resistant a filopodium/microspike is to bending sideways along its axis. Note that the parameter does not consider potential binding events between individual filaments or between filaments and other proteins, which might occur *in situ*. *For a more detailed mathematical description, see methods section.* | nm<sup>2</sup> |
| Parameter set describing entire network structures of randomly distributed filaments<br>(e.g. lamellipodia) |  |  |
| Parameter name | Parameter short description and biological relevance | Unit |
| Height (of structure) | Average height of the structure. It is averaged for 50 equidistant cross sections, considering the lowest and highest Z-coordinate points in each cross-section. | nm |

**Table 2.**
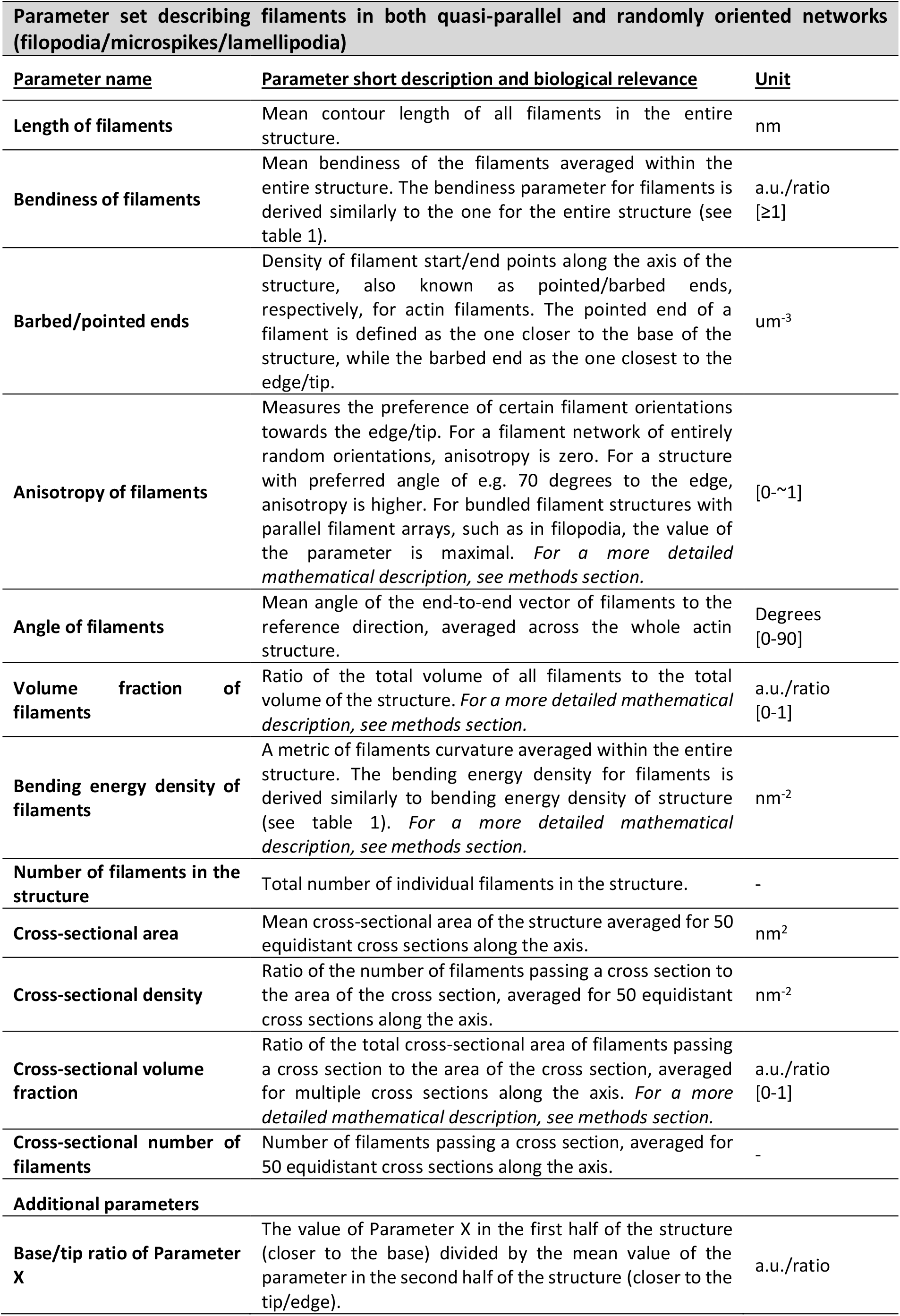
Description of parameters included in the computational toolbox, valid for both quasi-parallel filament arrays and dendritic networks.

### 3) Data curation

The low SNR and missing wedge effects in cryo-ET data often cause unwanted artifacts that, upon deriving coordinate files, result in false-positive filament tracking (Fig. 1C, Suppl. Fig. 2A). In order to reduce such false-positive information in downstream analysis, we implemented data cleaning and curation options to remove vector data of unspecific structures and background. Specifically, we implemented filtering of data files by custom ranges for filament length, angular distribution, or bendiness (Suppl. Fig-2A). The results of the cleaning steps can be fed into the visualization module integrated in the toolbox to receive feedback upon testing various parameters. Since the input and output format of the cleaned coordinates is also compatible with IMOD, an iterative manual manipulation of model files or cleaning of individual artifacts in IMOD and data analysis within the MATLAB-based toolbox is possible. To allow comparison of datasets acquired with different pixel sizes or fields of view (FOV) we have included an option to define pixel size and re-scale the dimensions of coordinate files in a semi-automated fashion. This enables the normalization of non-uniform datasets to compare differently acquired experimental groups (Suppl. Fig. 2B).

### 4) Simplified data interpretation and classification

We have integrated a GUI-based data visualization module, which works seamlessly with the output of the analysis scripts (Fig. 1D, Suppl. Fig-3A). It allows to review the quality of processed data, using various instruments, such as color-coding of filaments by customized parameter ranges (Suppl. Fig-3B), displaying cross sections along the axis, as well as overlaying 3D objects to extract representative images containing sufficient visual information (Suppl. Fig-3C). We facilitate the display of data and group comparisons by allowing to select the desired outcome through the user interface. Experimental groups can easily be assigned, compared visually by multiple types of readily available graphs, correlated to each other or classified via PCA analysis (Fig. 1E). Output of the analysis is also saved in .xls-files to allow a straightforward extraction of raw parameter values for various statistical tests or to feed them in other software workflows.

### Data analysis with the computational toolbox

In order to demonstrate the potential of the computational toolbox and its ability to dissect ultrastructural data and quantify differences between experimental groups, we compared distinctly organized branched networks or bundled arrays of filaments in vitreously frozen B16-F1 melanoma cells (Fig. 2A and B; Fig. 3). To this end we acquired cryo-electron tomograms of B16-F1 melanoma cells under different conditions. Cells were fixed and extracted as described previously (*10*) in order to preserve lamellipodia, filopodia and microspikes, while at the same time enhancing contrast due to the removal of membrane and cytosolic proteins. Filament coordinates were derived upon vitrification with either the filament segmentation package in the Amira-Avizo software package or a combination of deep-learning with the YAPiC software-based segmentation of filaments, followed by filament tracking in MATLAB scripts using the localized Radon transform (*15*). Both approaches can result in similar outcomes (Suppl. Fig-1). However, since obtaining filament coordinate information in Amira-Avizo required less manual user-defined parameter testing, increased throughput and also resulted in higher filament density, we decided to perform the remaining analysis presented in this manuscript using filament coordinates derived from Amira-Avizo. We note that segmentation using a convolutional neural network (CNN) like YAPiC can be applied in combination with any filament tracking approach.

**Figure 2.**
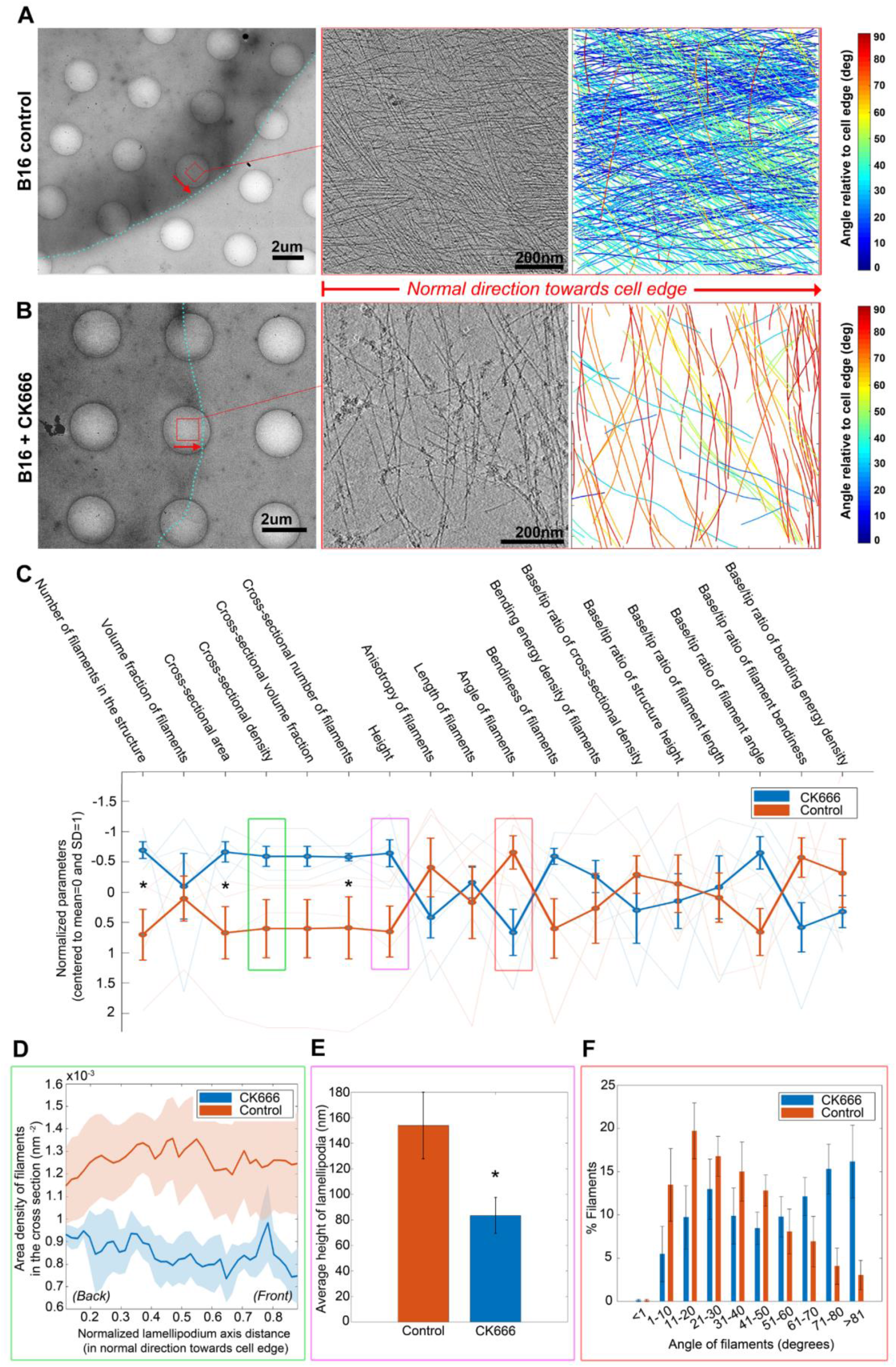
Analysis of lamellipodial networks. (**A-B**) Analysis of filament networks in cryo-electron tomograms of lamellipodia of **(A)** B16-F1 mouse melanoma cells treated with DMSO control or **(B)** treated with 210uM of the Arp2/3 inhibitor CK666. Left panels: Low-magnification overview images showing the cell periphery of the respective cells. The cell edges are annotated by a cyan dotted line. Middle panels: Summed 10 computational slices through bin8-tomograms of B16 lamellipodia. Their position at the cell periphery is annotated by red boxes in the left panel. Right panels: Visual output generated by the computational toolbox, color coded for angular distribution relative to the normal direction of the cell edge. Red arrows indicate the orientation of the axis towards the lamellipodial edge, i.e. normal direction. Scale bar sizes are annotated in the figure. **(C)** Normalized quantitative values of multiple parameters plotted linearly in one graph can reveal differences in ultrastructural characteristics between experimental treatments. Thick lines indicate the averaged values for all data files in a group, while faint lines the averaged (when appropriate) values of individual data files for every parameter. Information on the individual parameters is given in tables 1 and 2. Please note the orientation of the Y-axis. (**D-F**) Parameters selected from the linear graph in panel (C) plotted individually. **(D)** Cross-sectional filament density along the lamellipodial axis in normal direction to the cell edge. The transparent outlines indicate standard deviation. **(E)** Average lamellipodium height. **(F)** Angular orientation of filaments in lamellipodia (bin number and size is customizable). All plot options are easily accessible via a GUI-based module. Statistical significance (paired t-test, p≤0.05) between experimental groups is marked with *. N of tomograms is 4 for both control and CK666 groups

**Figure 3.**
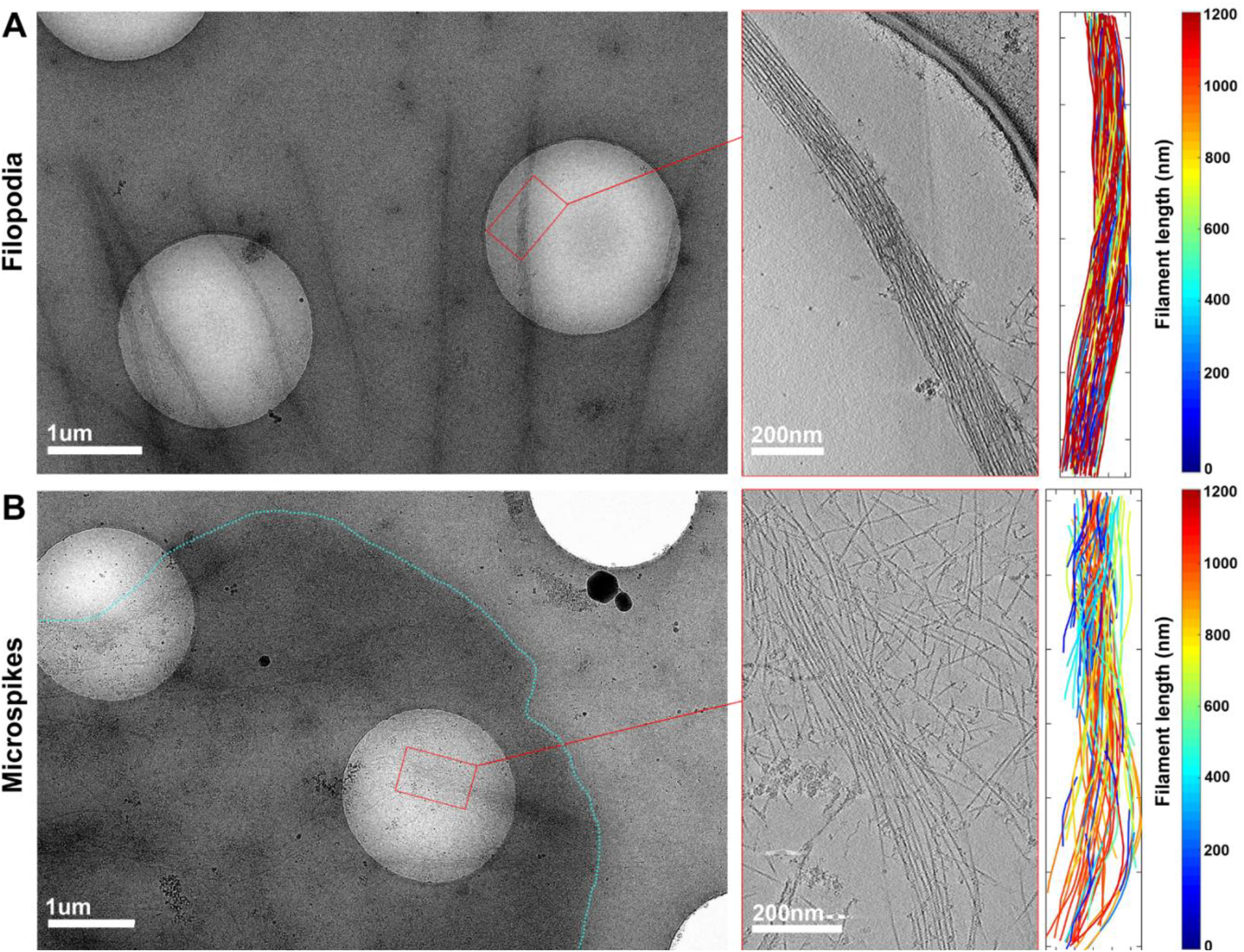
Representative EM micrographs of filopodia and microspikes used for analysis. In order to demonstrate the feasibility of the computational toolbox workflow in distinguishing quantitative ultrastructural differences between experimental groups containing quasi-parallel filaments, we compared **(A)** filopodia protruding beyond the cell edge to **(B)** posterior regions of microspikes embedded within lamellipodia. Filaments belonging to the lamellipodial networks were manually removed with IMOD. In (A) and (B), left panels show medium magnification maps of the cell periphery. Cyan dotted line indicates the cell edge. Analysed regions are highlighted with red rectangles; middle panels show a representative tomogram slice of the analyzed region; right panels show the visual output generated by the computational toolbox, color coded for filament length.

### Analysis of branched actin networks

The Arp2/3 complex is an integral component in dendritic actin networks. It binds to preexisting actin (mother) filaments, promotes the nucleation of new (daughter) filaments and thereby forms characteristic branch junctions, which link mother and daughter filaments (*23*). We analyzed either untreated (DMSO-control) B16-F1 cells (Fig. 2A, Suppl. Fig-4A) or B16-F1 cells treated with the Arp2/3 complex inhibitor CK666 (∼10min; 210uM concentration) (Fig. 2B, Suppl. Fig-4B) to compare branched filament networks with different architecture. CK666 binds to the Arp2/3 complex and inhibits actin filament nucleation by stabilizing the inactive state of the complex, thus also inhibiting dendritic actin network formation (*24*). As reported previously, CK666 treatment led to rearrangement of filaments, manifested by changes in their angular distribution and density in comparison to the untreated cells (*25*). While this is already discernible in the tomographic data, it becomes even more evident when the filament tracks are plotted and colored by ranges of angular distribution relative to the cell edge using our computational toolbox color-coding option (Suppl. Fig-4B). For straightforward comparison of datasets, our toolbox uses a linear visualization plot for all parameters describing filamentous networks/lamellipodia (Fig. 2C, Tables 1 and 2). This type of graph provides a convenient and fast approach to identify the key differences between experimental groups, which can then be analysed in detail with more specialized visualization options. For instance, filament density between control and CK666 treated cells can not only be averaged, but also traced along the axis of the entire structure (Fig. 2D) to identify potential differences between front and back regions of the structure. Other parameters, such as average lamellipodium height, are easily discerned by plotting them in a bar chart (Fig. 2E). Histogram plots can be used to compare the distribution of parameter values between experimental groups. In the presented case, this analysis confirms the visual impression of an increased fraction of filaments in CK666-treated cells, running in angles of >60 degrees to the cell edge, relative to control cells (Fig. 2F). Similarly, all parameters shown in Fig. 2C, can be displayed with various plots in order to separate filament populations in bins of custom size, discover potential differences in their values along the axis between two or more experimental conditions, find correlations or categorize data of sufficiently large size by e.g. PCA analysis (see Fig. 1E for example).

In addition, our toolbox allows the analysis of the distribution of filament start and end points along the structure axis. This type of analysis could reveal potential differences in the density of actin filaments pointed/barbed ends in back vs. middle vs. front regions of lamellipodia for each experimental condition, assuming that the pointed end of a filament is the one closer to the base and the barbed end – the one close to the edge/tip of the structure. (Suppl. Fig-5). An important consideration is the potential accumulation of these ends on the edges of cropped areas (or at the boundaries of the tomograms themselves) and hence the avoidance of including false-positives into the final analysis (Suppl. Fig-5A). We address this by allowing the user to set boundaries for selecting pointed/barbed ends lying in defined sections along the axis of the structure (i.e. away from the edge of the selected area) (Suppl. Fig-5B) or within certain distance ranges from the base or away from the tip of the structure.

### Analysis of bundled actin filament arrays

We used our toolbox for examining ultrastructural characteristics of bundled filament structures, and compared protruding filopodia with posterior regions of non-protruding microspikes (Fig. 3). Posterior regions show less uniform arrangement of filaments and are often diverging or splayed apart (as previously reported in (*26*)). On the contrary, protruding filopodia are characterized by more tightly bundled filaments (Suppl. Fig-6). Similar to the above described approach for analysis of branched networks, the toolbox enables displaying all parameters associated with bundled filament arrays in a linear plot (Fig. 4A), where multiple ultrastructural differences between filopodia and microspikes are immediately identifiable. Several of these parameters differ from the parameters shown for filament networks, such as lamellipodia, accounting for the bundle architecture. Also, in this case, individual parameters of interest can be plotted with other graph types in order to derive more information. For instance, visualizing the cross-sectional circularity parameter shows a clear reduction in back regions of microspikes, as opposed to their tip regions or to filopodia, likely indicative of less tightly bundled and irregular filament arrangement towards the back of the microspikes (Fig 4B). This corresponds to reduced values for the base/tip ratio of filament cross-sectional density and filament numbers in microspikes, as compared to filopodia (Fig. 4A). Differences in filament spatial arrangement and architecture between filopodia and microspikes are also evident when comparing their angles relative to the axis (Fig. 4C), as well as filament bendiness (Fig. 4D). The extracted quantifications clearly show that filaments in microspikes, as opposed to those in filopodia, are running at higher angles relative to the axis and are more bent, especially in the base of the structure. We implemented an alternative approach for spatial comparison of filament regularity in bundled structures by plotting angles and interfilament distances between filament pairs in cross-sections, based on work by Jasnin et al. (*12*) (Fig. 4 E,G). This allows identifying the abundance of parallel, regularly arranged filaments within the whole filament population. The presence of such filaments is characteristic of tightly bundled protruding filopodia, where an accumulation of parallel filaments separated by ∼10nm of interfilament distance is clearly evident (Fig. 4F, marked with white oval shape). Such arrangements are less abundant in the microspikes of our dataset (Fig. 4H). Similarly to the analysis of branched networks, all parameters shown in the linear comparison plot in Fig. 4A can be further analyzed using a large number of plots or using the visualization module of our toolbox to display cross-sections of structures at user-defined positions or characteristics (Suppl. Fig-3A).

**Figure 4.**
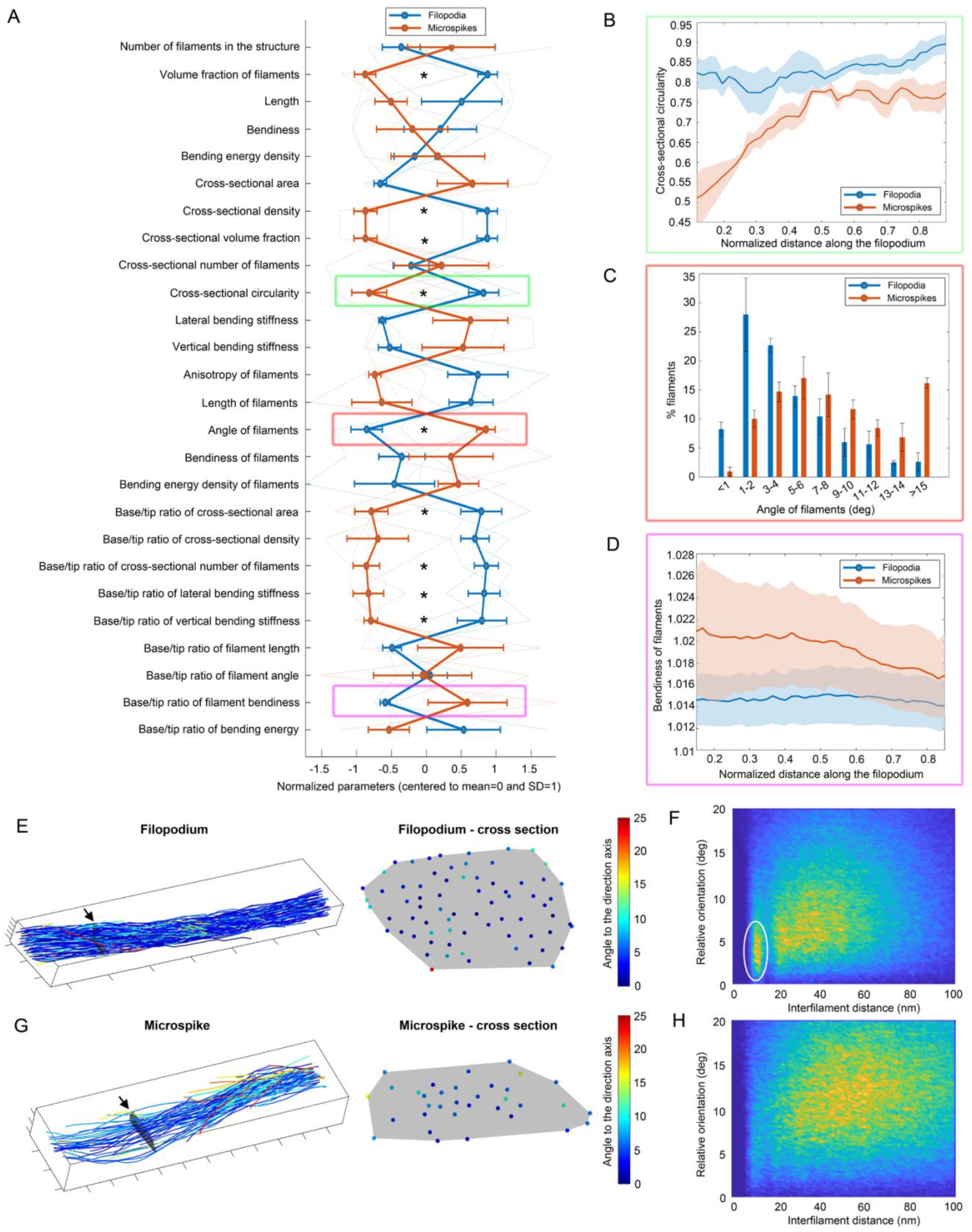
Analysis of bundled filament arrays. **(A)** Plotting normalized values of multiple parameters in one graph allows to reveal differences in ultrastructural characteristics of distinct structures. We compared a list of ultrastructural parameters between protruding filopodia and posterior microspikes. Individual parameters have been selected for more detailed comparison. These include: **(B)** average cross-sectional circularity of filopodia/microspikes along the axis (the transparent outlines indicate standard deviation), **(C)** angular orientation of filaments in each structure displayed with a histogram plot of customizable bin numbers and step sizes and **(D)** local bendiness of filaments along the axis of the structure. The computational toolbox also allows the visualization and extraction of quantitative information on the spatial organization of filaments relative to their neighbors. The integrated visual module was used to first display and example of a **(E)** protruding filopodium and **(G)** posterior microspike, where left panels display the analyzed structure with black arrows indicating the location of cross-sectional segments along the axis, and right panels show the cross-sectional distribution of individual filaments color-coded by their local angular orientation to the axis. **(F, H)** Relating distances between filament pairs (in nm) to their relative local orientations (in degrees) demonstrates the presence of a higher number of tightly bundled and parallel oriented filaments within filopodia (indicated with white oval in F) compared to posterior microspikes (H). All plot options are easily accessible via the GUI-based module. Statistical significance (paired t-test, p≤0.05) between experimental groups is marked with *. The number of tomograms is 3 for both filopodia and microspike groups.

## Conclusion

Here we introduce a MATLAB-based computational toolbox, which facilitates the processing and analysis of filament-rich ultrastructural data extracted from cryo-electron tomograms. As a proof-of-principle, we have analyzed a relatively small sample size and compared parameters and experimental samples with obvious differences in filaments distribution. Within this manuscript we did not intend to reveal new biological insights into the actin network architecture, but rather showcase the functionalities of the introduced toolbox. We expect that with increased throughput in data acquisition (*4*–*6*), large datasets for a variety of samples can be acquired in short time, further highlighting the importance to develop ease-of-use tools allowing efficient analysis of the wealth of biological data contained within cellular cryo-electron tomograms. Indeed, we believe that the real power of the toolbox comes with the time-efficient analysis of large datasets, and will allow the detection of subtle ultrastructural differences between experimental conditions, e.g., when comparing multiple genetic knockout clones with mild phenotypes.

In such a case, purely visual comparisons, or even smaller scale analysis, might prove to be inconclusive. The wealth of data can be useful for a better understanding of the role of proteins contributing to a given network, or can supplement or enable mathematical modeling approaches of network initiation and maintenance (*27, 28*).

While we developed the toolbox with an emphasis on cellular actin networks and actin-rich cell peripheral structures, it can in principle be used for the analysis of virtually any filamentous network in ET data (or other imaging data) belonging to two different types of ultrastructural assemblies: filament networks (such as branched networks within lamellipodia); and filament architectures, which are aligned in a quasi-parallel fashion (such as filopodia or microspikes). There are numerous examples for biological filamentous assemblies for which this analysis is expected to be applicable, e.g., other cytoskeletal elements, such as microtubules or intermediate filaments, and extracellular networks composed of fibrilous components, such as collagen or fibronectin. The modality of the toolbox allows the implementation of additional features and parameters in the future in order to increase its adaptability for more specialized projects or for investigating other structural configurations.

## Acknowledgements

This research was supported by the Scientific Service Units (SSUs) of IST Austria through resources provided by Scientific Computing (SciComp), the Life Science Facility (LSF), the BioImaging Facility (BIF), and the Electron Microscopy Facility (EMF). We also thank Victor-Valentin Hodirnau for help with cryo-ET data acquisition. The authors acknowledge support from IST Austria and from the Austrian Science Fund (FWF): M02495 to G.D. and Austrian Science Fund (FWF): P33367 to F.K.M.S.

## Author contributions

Georgi Dimchev: Conceptualization, Methodology, Software, Validation, Formal analysis, Investigation, Data Curation, Writing - Original Draft, Writing - Review & Editing, Visualization, Project administration, Funding acquisition. Behnam Amiri: Conceptualization, Methodology, Software, Data Curation, Writing - Original Draft, Visualization. Florian Fäßler: Methodology, Software, Validation, Investigation, Writing - Review & Editing. Martin Falcke: Supervision, Writing - Review & Editing, Funding acquisition. Florian Schur: Conceptualization, Validation, Writing - Original Draft, Writing - Review & Editing, Visualization, Supervision, Project administration, Funding acquisition.

## Code availability

The MATLAB toolbox scripts including a detailed documentation on their usage and example data for processing are available for download via https://schurlab.ist.ac.at/downloads/

## Materials and Methods

### Cell culture

B16-F1 mouse melanoma cells were kindly provided by Klemens Rottner (Technical University Braunschweig, Helmholtz Centre for Infection Research). Cells were grown at 37°C and 5% CO_2_ and cultured in Dulbecco’s modified Eagle’s medium (DMEM GlutaMAX, ThermoFischer Scientific, #31966047), supplemented with 10% (v/v) fetal bovine serum (ThermoFischer Scientific, #10270106) and 1% (v/v) penicillin-streptomycin (Thermo Fischer Scientific, #15070063).

### Cryo-ET sample preparation and inhibitor treatments

B16-F1 cells were cultured as described above and seeded onto 200 mesh gold holey carbon grids (R2/2-2C; Quantifoil Micro Tools). Prior to cell seeding, the grids were placed onto a piece of parafilm sheet, firmly attached to the bottom of a 6-well flat bottom dish, and coated for 1hr RT with 25 μg/ml laminin (Sigma, L2020) diluted in a buffer containing 150 mM NaCl, 50 mM Tris, pH 7.5. Grids were gently washed with PBS and cell suspension was pipetted into the well in a slow drop-wise fashion to avoid flipping of the grids.

For CK666 inhibitor treatment, the medium of adherent cells grown overnight onto the EM grids was gently replaced with growth medium supplemented with either 210µM CK666 (Sigma Aldrich, #SML0006) or an equivalent amount of DMSO. An incubation time of 10min with the inhibitor was chosen in order to allow CK666 to induce defects in the organization of the actin filaments network in lamellipodia, while not causing the complete retraction of the structure.

Following either overnight growth (for filopodia/microspikes) or overnight growth followed by a 10min treatment with DMSO/CK666 inhibitor, cells were extracted and fixed as previously described (*19*). In brief, grids were incubated for 1min RT in a drop of cytoskeleton buffer (10mM MES, 150mM NaCl, 5mM EGTA, 5mM glucose and 5mM MgCl2, pH6.2) supplemented with 0.75% Triton X-100 (Sigma-Aldrich, #T8787), 0.25% glutaraldehyde (Electron Microscopy Services, #E16220) and 0.1μg/mL phalloidin (Sigma-Aldrich, #P2141). Fixation was subsequently performed for 15 minutes at RT by placing the grids in a drop of cytoskeleton buffer containing 2% glutaraldehyde and 1μg/mL phalloidin.

Following extraction and fixation, grids were subjected to back-side blotting (3sec blot time) and vitrification using a Leica GP2 plunger equipped with a blotting detection sensor and incubation chamber maintaining an environment of 21°C and 90% humidity (Leica Microsystems). Grids were placed into the GP2 incubation chamber and excessive liquid was manually removed with a piece of filter paper by gently touching the side of the grid. Prior to blotting and plunging into liquid Ethane (−185°C), 3μl of a solution of 10nm colloidal gold (AURION Immuno Gold Reagents & Accessories, Netherlands) coated with BSA in PBS was added onto the grids. Samples were placed in liquid nitrogen storage until imaging.

### EM Data acquisition

Tilt-series were either acquired on a Thermo Scientific 300kV Titan Krios G3i TEM equipped with a with a BioQuantum post-column energy filter and a K3 camera (Gatan) or on a Thermo Scientific 200kV Glacios Cryo-TEM equipped with Falcon 3EC camera. Both microscopes were aligned and operated using the SerialEM package (*29*).

For data acquired on both microscope systems, the workflow included acquisition of low- and medium-magnification montages for defining regions of interest, followed by high-resolution data acquisition with varying magnification settings and pixel sizes for the different experimental groups (described below).

All filopodia and microspikes data were acquired on a Titan Krios G3i TEM with a total electron dose of ∼180e^-^/px, a tilt range of -62/+62 degrees with 2-degree steps and a defocus of ∼-3um. Two filopodia and three microspikes were acquired with a pixel size of 2.137 Å(magnification of 42,000x), while one filopodium was acquired with pixel size of 2.676 Å and magnification of 33,000x.

Two of the untreated lamellipodia were acquired on Titan Krios Krios G3i TEM with a total electron dose of ∼ 180e-/px, pixel size 2.137 Å (magnification 42,000x) and defocus of ∼-4um.

Two of the untreated lamellipodia and all lamellipodia treated with CK666 were acquired on a Glacios TEM with a total electron dose of ∼150e^-^/px, tilt series of -62/+62 degrees with 2-degree steps, defocus of ∼-3um and pixel size of 3.24 Å (magnification of 45,000x).

### EM data processing and extraction of coordinate files

Pre-processing of acquired tilt series (tilt stack sorting, removal of bad tilts, exposure filtering) was performed with the MATLAB-based Tomoman package(*30*). Tomogram reconstruction from the filtered tilt series was performed with the IMOD/Etomo software package.

As illustrated on Suppl. Fig-1, two different approaches were applied for actin filament vectorization and extraction of filament coordinates from reconstructed tomograms. The first approach involves using tomograms as input for training neuronal networks via the interactive learning and segmentation toolkit Ilastik (*31*) and the YAPiC pixel classifier (*22*). Actin filaments and background were manually annotated in Ilastik and the the Ilastik-derived .ilp project files were processed in YAPiC to generate model files, i.e. trained neuronal network instructions for automated segmentation of filaments in a larger dataset. Separate trainings were performed for lamellipodial networks (with bin8 tomograms) and filopodia/microspikes (with bin4 tomograms). YAPiC-derived model files were used by the same software to generate binary prediction files from reconstructed tomograms, distinguishing between filaments and background.

YAPiC-derived prediction files were processed with MATLAB scripts using the localized radon transform (*15*) allowing the extraction of files containing XYZ coordinates of points assigned to individual filaments. Cleaning of false-positive filaments was additionally performed via a custom-made Python script eliminating filament pairs within a defined proximity (in pixels) to each other. Another approach involved processing reconstructed tomograms with the Amira-Avizo software package, using the “Cylindrical correlation” and “Trace correlation lines” modules. The following parameter values for the Cylindrical correlation module were set for raw (i.e. header-containing) tomogram .rec files of bin8: Cylinder Length=500; Angular Sampling=5; Mask Cylinder Radius=45; Outer Cylinder Radius=35; Inner Cylinder radius=0 (all units are in Å). The following parameter values were set for the Trace Correlation Lines module: Minimum Seed correlation (tomogram dependent, varying between 80-120); Minimum Continuation Quality=100; Direction Coefficient=0.3; Minimum Distance=70; Minimum Length=350; Search Cone Length=500; Search Cone Angle=37; Search Cone Minimum Step Size(%)=10. Segments and point coordinates were extracted as separate excel sheets from Amira-Avizo and reformatted with a custom-made MATLAB-script (“amira_reformat_to_coordinates.m” script provided together with the computational toolbox) in order to obtain a single file per tomogram containing XYZ coordinates of points assigned to each individual filament.

### Data pre-analysis cleaning and processing

Prior to analysis of data files with the computational toolbox, cleaning of unspecific background and false positives was performed. Unspecific background was removed by using the filtering scripts included in the “Supplemental scripts” folder of the toolbox, excluding all filaments with a length of less than 100nm and an angle of less than 75 degrees in Z axis (as illustrated in Suppl. Fig-2A). Individual unfiltered filaments, as well as filaments belonging to lamellipodial networks around microspikes, were manually removed with the IMOD software. The “point2model” and “model2point –c” functions were used to re-format respectively .txt coordinate files into IMOD-compatible .mod files or vice versa. For lamellipodia, area of all data files was normalized to 800×800nm in XY, by using the cropping script provided in the “Supplemental_scripts” folder of the MATLAB toolbox (see Suppl. Fig-2B).

### Software packages used for manuscript assembly and figures preparation

Coding of the computational toolbox was performed in MATLAB (The MathWorks Inc.). All statistics were performed with the SigmaPlot software (Systat Software Inc.). Figures assembly and preparation was performed with Adobe Photoshop and Adobe Illustrator (Adobe Inc.).

### Description of ultrastructural parameters

Since some basic definitions are used repetitively in the description of parameters, we introduce these definitions first.

### Reference direction

Many of the ultrastructural metrics/parameters described later are dependent on a reference direction. In filamentous networks, such as lamellipodia, it is defined as the direction of a vector pointing towards the leading edge (identical to the axis of lamellipodium or normal direction towards the cell edge). In quasi-parallel filamentous arrays, such as filopodia, it is defined as the direction of a vector starting from the base of the structure to its tip (see axis labeled with “X” in Suppl. Fig-7A). Note that this vector is not identical to the axis of the filopodium, which may be a curved line.

### Axis and cross sections

In the filament networks (e.g. lamellipodia), the axis is a vector pointing towards the leading edge. In the quasi-parallel arrays (e.g. filopodia/microspikes), the axis may also be a curved line from the base to the tip, which follows the curve of the filopodium. In the latter case, the axis is determined using a second order polynomial fitting in the X-Y plane on the data points of all the filaments in the structure. A cross section is defined at a point on the axis, as the plane perpendicular to the direction of the local tangent at that point.

### Global and cross-sectional frames of reference

We define a global frame of reference based on the reference direction. The basis of this frame is defined as follows (see also Suppl. Fig-7A): X points towards the reference direction; Z points in the direction perpendicular to the cell plane (the X-Y plane on which the structure is lying). Note that it is assumed that the filament points in z indicate a coordinate perpendicular to the cell plane. Finally, Y is defined as the cross-product of Z and X. Similarly, we define a local cross-sectional frame of reference at every cross section (See Suppl. Fig-7B). The basis of the local frame of reference are: x’ is the local tangent vector of the axis; z’ points in the direction perpendicular to the cell plane (similar to Z), and y’ is the cross-product of z’ and x’. Origin of this local cross-sectional frame of reference is at the center of the mass of the cross section.

### Bending energy density (of entire filopodia structures)

Bending energy density for a contour (in this case the axis of the structure) is defined as the normalized sum of the squared local curvature on the contour, i.e.

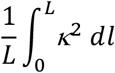

where *κ* is the local curvature at any point on the contour; *l* is the distance along the contour; *L* is the contour length.

### Volume fraction of filaments

Ratio of the total volume of all filaments to the total volume of the structure. Each filament is assumed to be a cylinder with the diameter *d* (default diameter is 7nm). The total volume of filaments is:

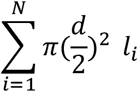

where *N* is the total number of filaments in the structure and is the length of i’th filament.

### Bending energy density of the filaments

Mean bending energy density of the filaments averaged in the whole actin structure. It is derived similarly to the Bending energy density for entire structures parameter, described above. However, as opposed to Bending energy of a structure, which is calculated based on the structure axis, Bending energy density of the filaments parameter is derived from the contour of individual filaments within the structure.

### Anisotropy of filaments

Mean squared deviation of the angular distribution of the filaments from the uniform angular distribution.

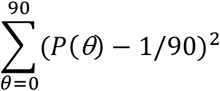

where *P(θ)* is the probability distribution function of filament angle; *θ* is the angle of filaments to the reference direction in X-Y plane.

### Cross-sectional volume fraction

Ratio of the total cross-sectional area of filaments passing a cross section to the area of the cross section, averaged for 50 equidistant cross sections along the axis. Cross section of each filament is assumed to be a circle with diameter *d* (default diameter is 7nm). Thus, the total cross-sectional area of filaments is *n*π(*d*/2) ^2^, where *n* is the number of filaments passing the cross section.

### Cross-sectional circularity (only for filopodia/microspikes)

Mean cross-sectional circularity of the filopodium averaged across 50 equidistant cross sections along the axis. Circularity is defined as the ratio of the cross-sectional area to the area of a circle with the same perimeter.

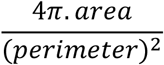

For a perfect circular cross section circularity equals 1. It is less than 1 for not circular cross sections.

### Lateral bending stiffness (only for filopodia/microspikes)

The moment of inertia of the filaments in a cross section with respect to the z’-axis, averaged across 50 equidistant cross sections along the axis. This parameter describes the resistance against lateral bending.

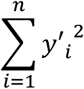

where n is the number of filaments passing the cross section; *y*′_*i*_ is distance of the filament *i* from the z’-axis (Suppl. Fig-7B).

### Vertical bending stiffness (only for filopodia/microspikes)

The moment of inertia of the filaments in a cross section with respect to y-axis, averaged across 50 equidistant cross sections along the axis. This parameter describes the resistance against vertical bending.

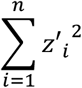

where n is the number of filaments passing the cross section; *z*′_*i*_ is distance of the filament from the y’-axis (Suppl. Fig-7B).

### Parameters describing properties along the axis (Derived from the Plots_Group_Cell_Filopodia/Lamellipodia.m script in the “Properties along axis” menu)

This script illustrates the variation of the properties of the structure along the axis, determined in 50 equidistant cross sections along the axis. To calculate the properties that are related to the filaments (length, angle, bendiness, and bending energy density) at a cross section, we average that property across all the filaments that are passing through the cross section.

### Parameters describing the spatial arrangement of filaments (Derived from the Plots_Group_Cell_Filopodia/Lamellipodia.m script in the “Configuration of filaments” menu)

We determine the relative distance and orientation of all filament pairs to describe their spatial organization within the structure. To calculate these parameters, a local normal plane at every point along a reference filament is defined (grey rectangle, Suppl. Fig-7C). This normal plane is perpendicular to the tangent vector of the reference filament (dark blue vector in Suppl. Fig-7C). The intersections of all other filaments with this plane are then determined. A relative orientation is defined as the angle between the tangent vector of each of these filaments at their intersection with the normal plane (light blue vectors in Suppl. Fig-7C) and the tangent vector of the reference filament at its intersection point. The interfilament distance is then defined as the distance between the intersection points of each filament and the reference filament with the normal plane. This procedure is repeated for 200 points along each filament and reiterated for every filament in the structure.

## Supplementary Information

**Suppl.Fig-S1.**
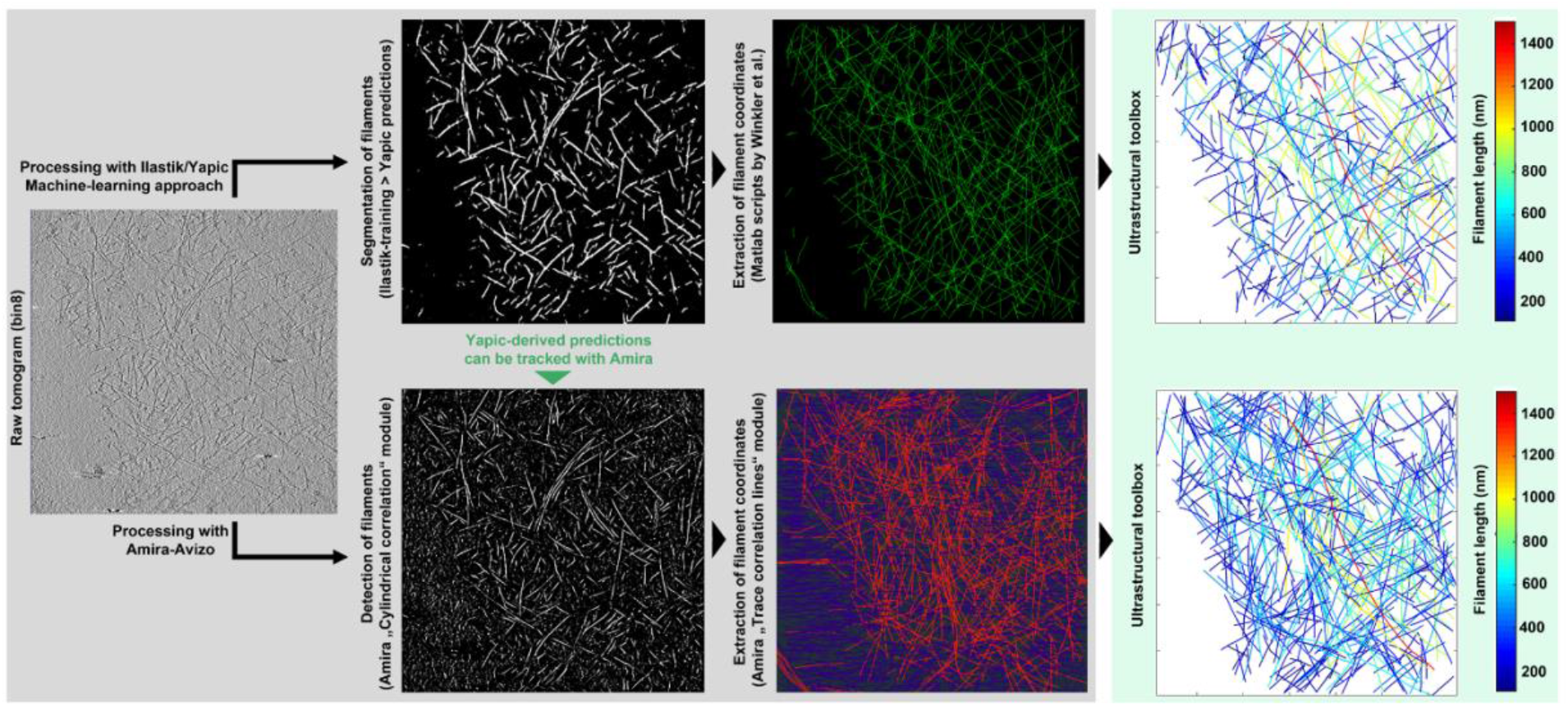
Workflow for extraction of filament coordinate data from cryo-electron tomograms. Here, we compare two alternatives, but not mutually exclusive approaches for filament vectorization and extraction of coordinate files. The first approach (upper row, left image) applies the interactive learning and segmentation toolkit Ilastik, in combination with the YAPiC pixel classifier, to train neuronal networks and generate binary image stacks facilitating the separation of filaments from background. Extraction of filament coordinates data from prediction files (middle panel) can then be performed by available MATLAB scripts (*15*). An alternative approach (bottom panels) applies the Amira-Avizo software package, where filaments are detected and separated from background via the “Cylindrical correlation” module (left panel) and traced via the “Trace correlation lines” module (middle panel). Notably, the YAPiC-derived prediction files can also be used with Amira-Avizo for downstream segmentation and extraction of filament coordinates information. Right images in both upper and lower panels indicate the visual output extracted from the computational toolbox, color-coded for filament length.

**Suppl.Fig-S2.**
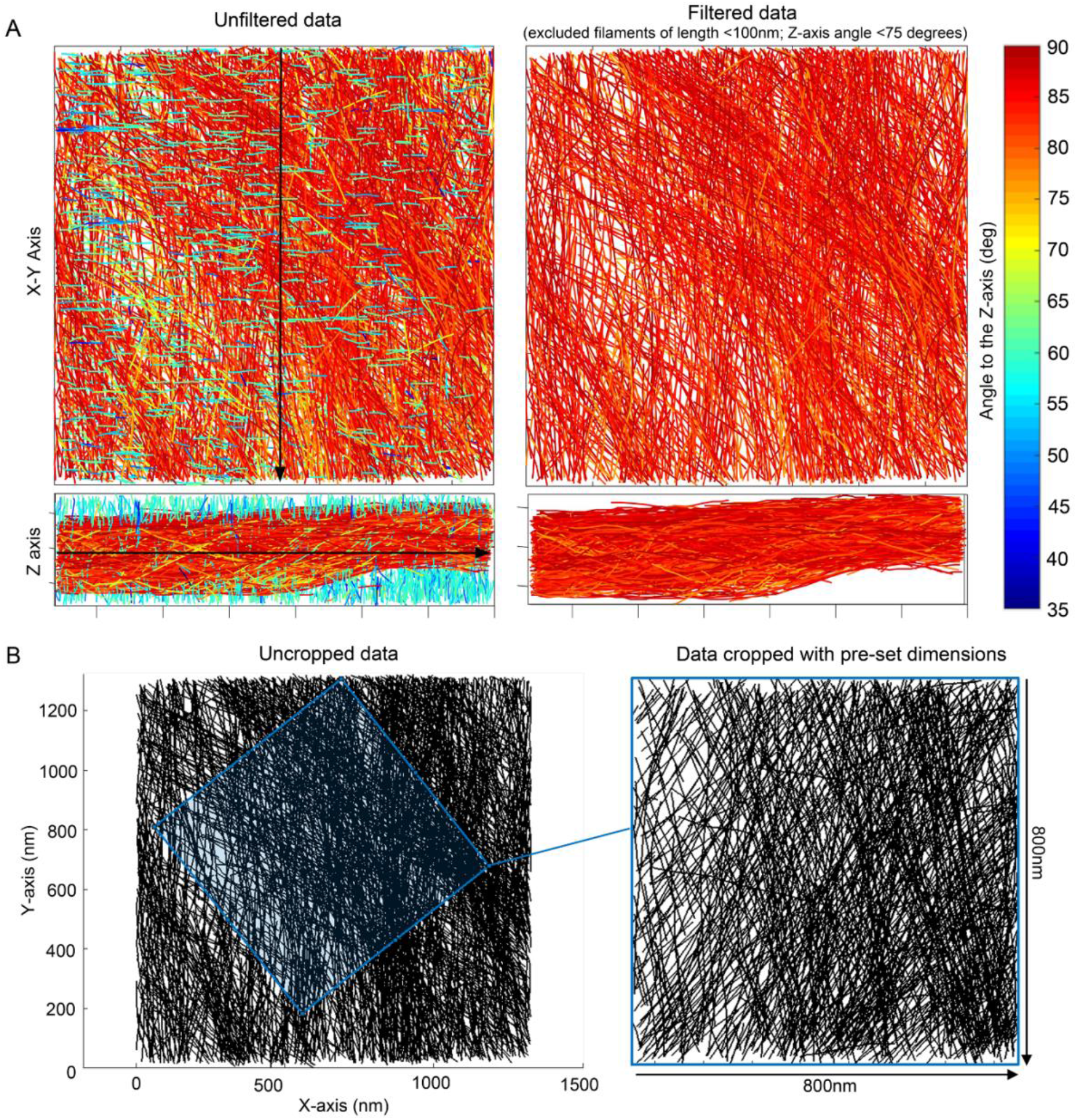
Data curation. **(A)** The toolbox allows removal of unspecific background and to select for analysis only filaments of specific characteristics, via filtering filaments in user-specified ranges for length, bendiness or angular distribution to X- or Z-axis. The example shows a filamentous network containing unspecific background, manifested as multiple short filaments running approximately perpendicular to the specified X-axis (green colored filaments in left panel). The filament network is shown before (left) and after (right) cleaning. The orientation of the X-axis is indicated with a black arrow. (**B)** Datasets containing filament coordinate files of non-uniform dimensions or pixel size, can be normalized for downstream analysis by a supplemental cropping script provided with the computational toolbox. An area of desired dimensions can be specified and manually positioned within the coordinate system of each data file (left panel) in order to obtain an output containing only the filament coordinates within the specified dimensions (right panel).

**Suppl.Fig-S3.**
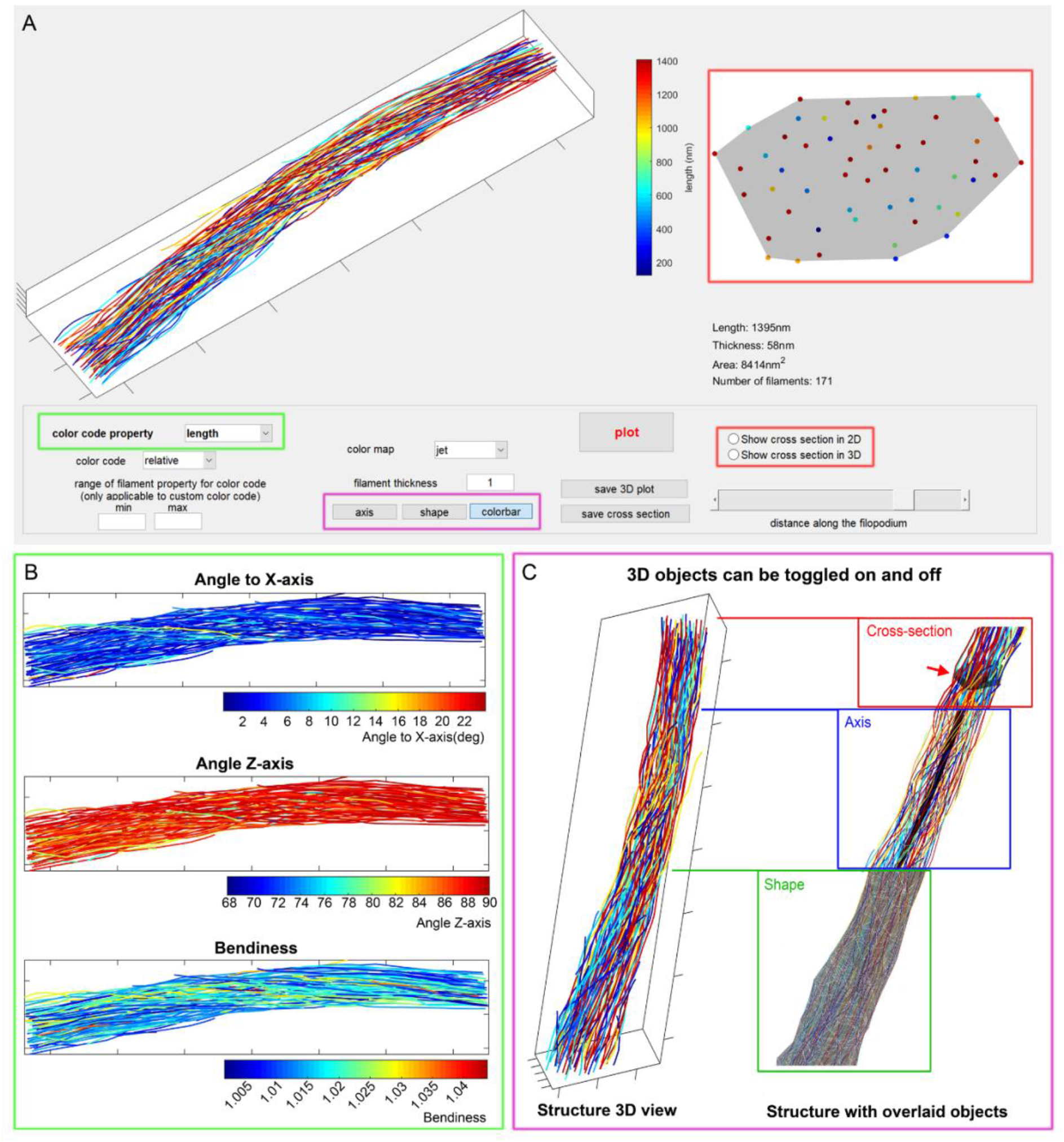
Visualization module. **(A-C)** The GUI-based 3D visualization module integrated in the computational toolbox allows for customized structure visualization. Filament color coding can be based on selectable ranges for length, bendiness or angular orientation (the green rectangle shows the selected option in panel **(A)** and three alternative output examples shown in panel **(B)**. Objects, such as axis, shape or color bar can be displayed together with the structure (purple rectangles in **(A)**) with respective examples in **(C)**. The module allows to visualize the cross-sectional orientation of filaments (red rectangles in **(A)**), with the position of the cross-section being adjustable along the axis of the structure (red arrow in C). More information on the structure such as length, thickness, area covered and number of filaments contained is also given below the cross-section visualization panel.

**Suppl.Fig-S4.**
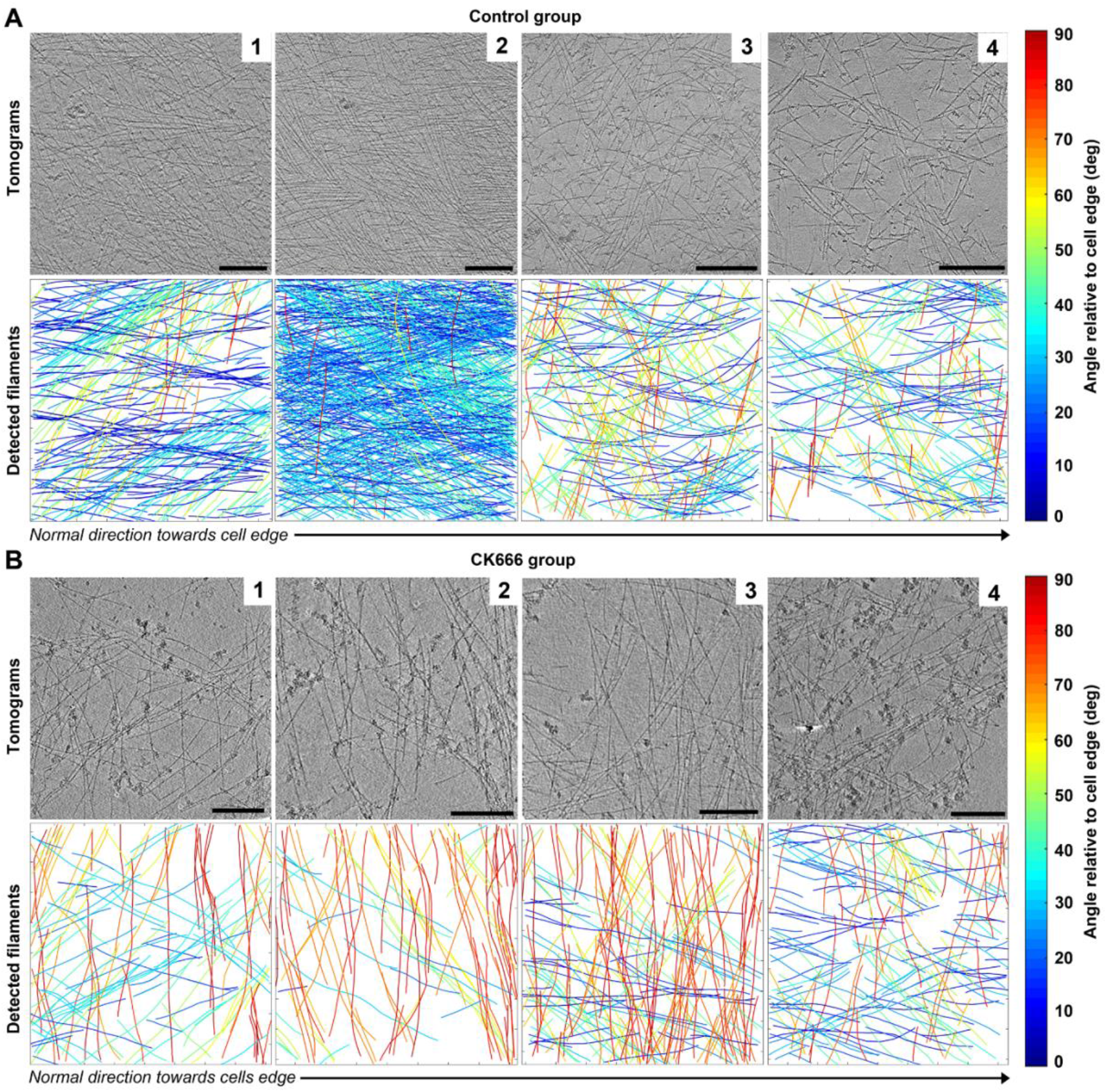
Visual gallery of lamellipodial dataset. A visual representation of all data files considered for quantitative analysis for **(A)** Lamellipodia of B16-F1 cells treated with DMSO control or **(B)** cells treated with Arp2/3 inhibitor CK666. Upper panels show 10 summed representative bin8-tomogram slices, bottom panels show computational toolbox-extracted visual output of analysed coordinate files, color-coded for angular distribution of filaments to the cell edge. Black arrows indicate the orientation of the axis towards the cells edge in normal direction. All scale bars correspond to a length of 200nm. Note that the second panels (from left) for each group are also displayed in Figure-2.

**Suppl.Fig-S5.**
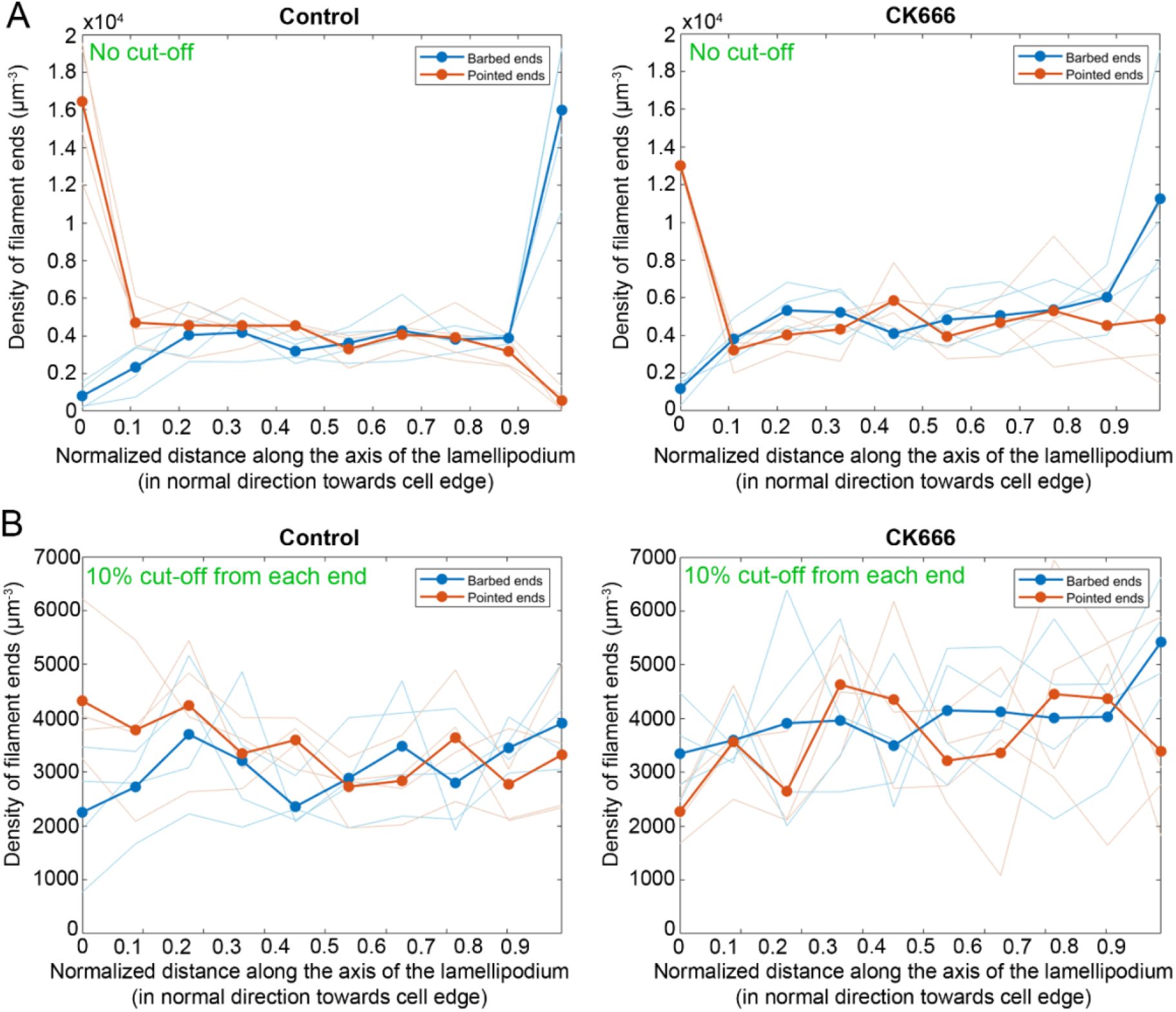
Distribution of filament barbed and pointed ends along the lamellipodial axis. (**A-B**) The distribution of barbed and pointed ends of actin filaments is shown for B16-F1 cells treated with DMSO control (left panels) or with Arp2/3 inhibitor CK666 (right panels). **(A)** No edge boundary cut-off was considered with obvious accumulation of pointed and barbed ends at the back and front regions of the structure, respectively. **(B)** Same representation of the distribution of filament barbed and pointed ends along the axis of the lamellipodia, but excluding those positioned within the first and last 10% of the axis length. For all panels, thick lines indicate the values of barbed/pointed ends averaged for all data files in a group, while faint lines indicate the averaged values for individual data files.

**Suppl.Fig-S6.**
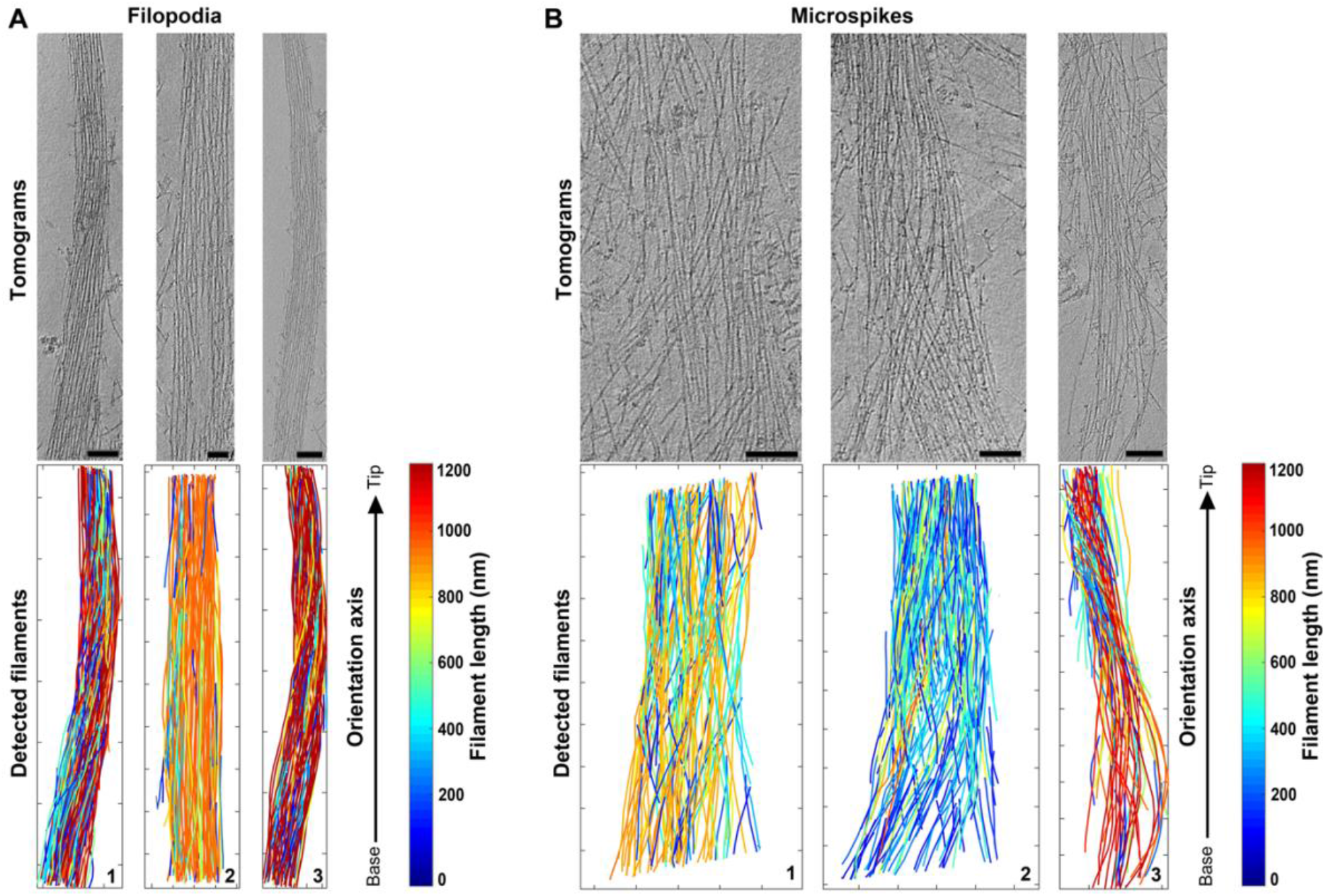
Visual gallery of filopodia/microspikes dataset. A visual representation of all data files considered for quantitative analysis for **(A)** filopodia or **(B)** microspikes. Upper panels show 10 summed representative bin8-tomogram slices, bottom panels show a typical visual output from the computational toolbox of analysed coordinate files, color-coded for filament length. Black arrows indicate the direction of the axis towards the tip of the structure. All scale bars correspond to a length of 100nm. Note that the 3rd panels (numbers are indicated in the figure) for each group are also displayed in Figure-3.

**Suppl.Fig-S7.**
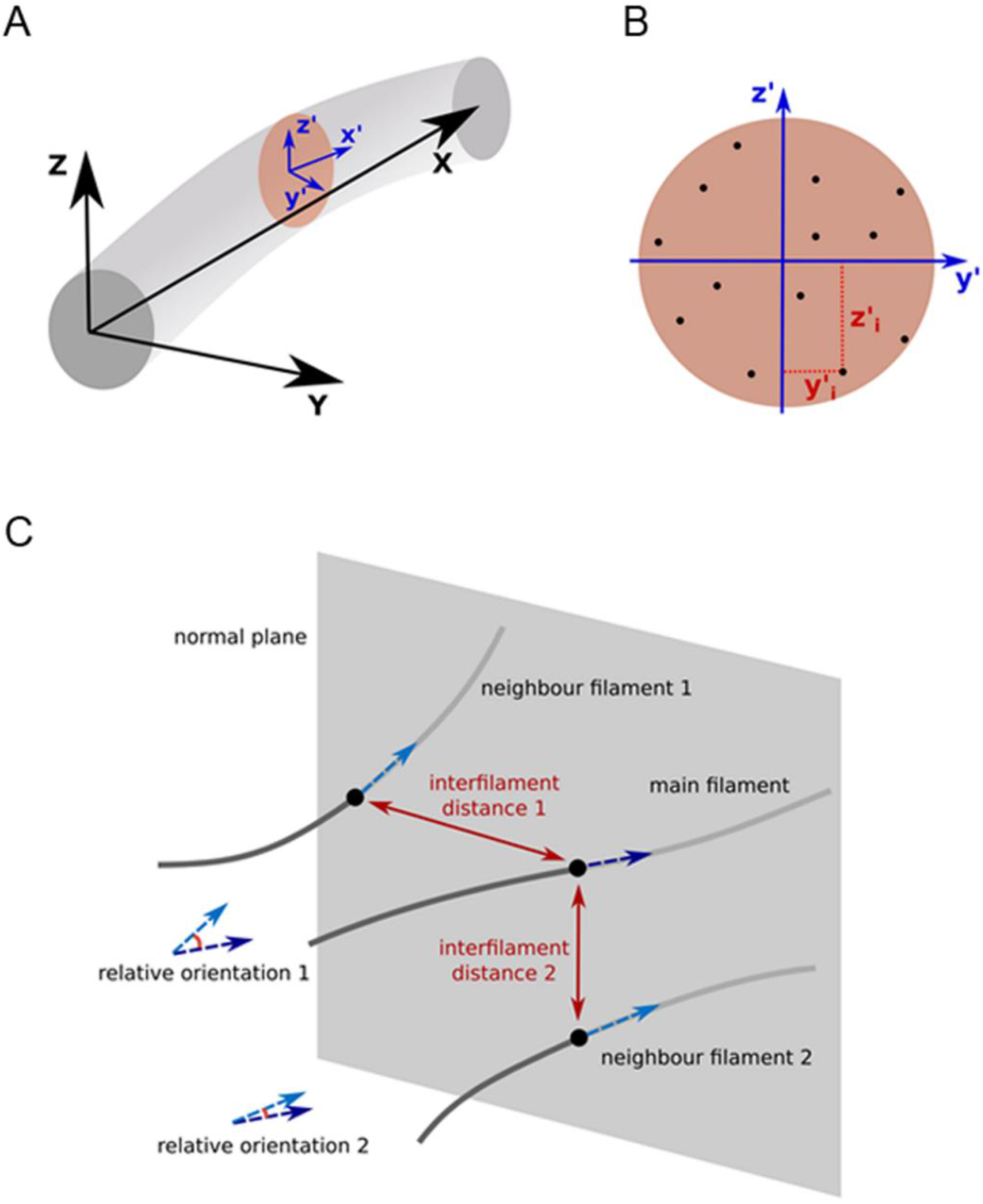
Visual representation of geometry-based definitions used for deriving ultrastructural parameters. The figure supplements understanding of the parameter description presented in the methods section. **(A)** Global frame of reference is shown with black letters and vectors. The local cross-sectional frame of reference is shown with blue letters and vectors for an arbitrary cross section along the filopodium. **(B)** An example of defining the coordinates of a filament in the local cross-sectional frame of reference. Black dots indicate individual filaments, intersecting with the cross section. **(C)** Interfilament distances and relative orientations at a certain point on a filament with respect to two neighbor filaments. Normal plane is a plane perpendicular to the main filament at an arbitrary point. Black dots show the intersections of filaments with the normal plane. Dashed vectors are the tangent vectors of the filaments at the intersection points.

